## Supplementary Information for "Pathogen transmission from vaccinated hosts can cause dose-dependent reduction in virulence"

The design and sampling protocol of the MD transmission experiment presented in the main manuscript builds on results from previous small-scale pilot experiments to establish (i) the appropriate contact duration between shedder and contact birds required for successful virus transmission, (ii) first estimates for the onset and duration of the infectious period of shedder birds and the impact of vaccination on it, and (iii) the appropriate time for collecting feather samples for detecting virus in contact birds infected by shedder birds. All pilot experiments were carried out in the same facilities as the main experiment (Avian Disease and Oncology Laboratory, ADOL, East Lansing, MI USA). Unless otherwise stated, the same experimental protocol was adopted for all experiments.

**(i) Determination of the duration of contact between shedder and contact birds required for successful virus transmission.** In this pilot experiment, contact birds came from the same chicken line as in the main experiment (maternal antibody negative 15I_5_ x 7_1_ F_1_), while shedders belonged to the inbred white leghorn chicken Lines 6_3_ (resistant to MD) and 7_2_ (susceptible to MD), all developed at the ADOL^1^. For each replicate, 28 newly hatched contact birds were placed with three-week old shedder birds from either line 6 or 7. Contacts were randomly allocated into four groups (hence 4x28=112 contact birds total). Subsequently, on each of hours 4, 8, 12, 24, 48, 96 and 168, four contact birds from each replicate were removed and placed together in an isolator (different isolator for each replicate, i.e., 4 birds per isolator). Isolated contacts were then monitored for 8 weeks and necropsied to determine MD status. Hence there were two independent groups of four contacts per shedder line per time period.

The proportion of contact individuals showing visible disease symptoms upon necropsy at 8 weeks post-contact was universally high in this experiment (Fig S1). Hence, it was concluded that 48 hours of contact is sufficient to establish MDV transmission from shedders with both high and low MD genetic resistance.

**Fig S1. Effects of exposure duration on contact bird Marek’s disease.** For each tested contact duration, the proportion of line 15I_5_ x 7_1_ F_1_ contact birds positive for Marek’s disease symptoms at necropsy, 8 weeks post-contact with inoculated unvaccinated “MD-resistant” line 6 (blue line) or “MD-susceptible” line 7 (red line) shedder birds.

**(ii) Establishing the onset and duration of the infectious period of vaccinated and sham-vaccinated MDV-infected shedder birds.** We carried out two replicates of experiments to examine whether vaccination affects the onset and duration of MDV transmission of infected shedder birds, and whether this is reflected by differences in shedder feather viral load over time. Groups of 3 shedder individuals of the same vaccination status were placed in contact with new independent groups of 10 contact birds every 2 days from 10-20 DPI, and recording of contact bird infection (qPCR at 14 DPC) and disease (necropsy at 8 weeks post-contact) status was carried out. The same inbred chicken line as the main experiments, 15I_5_ x 7_1_ F_1_, was used for both shedders and contacts, and protocols otherwise matched those of the main experiment. Shedder FVL was recorded at the beginning of each contact period, as well as at 7 DPI and weekly from days (DPI) 21-56 (see Fig S2).

Inspection of FVL profiles fitted to the repeated FVL measures of vaccinated and non-vaccinated shedders (Fig S2) showed that vaccinated shedders had consistently lower FVL than sham-vaccinated shedders and that both types of birds experienced an initial sharp rise in FVL. However, in sham-vaccinated shedders this initial sharp rise started to plateau at around 11-12 DPI, whereas FVL reached a peak at around 20 DPI in vaccinated shedders (Fig S2). The profiles suggest that any FVL-mediated effects of shedder vaccination on contacts should differ more at the earlier DPI. Given these results, DPI 13 and 20 were chosen as the most informative time points to investigate vaccination effects on MDV transmission and subsequent disease progression in contact birds in the main experiment.

**Fig S2. Feather viral load over time for shedder birds.** Vaccinated (blue) and sham-vaccinated (red) shedders, with maximum likelihood broken stick regression lines indicating lower viral load and a later breakpoint in viral load over time for vaccinated shedders. Open circles = replicate 1, crosses = replicate 2. The shaded area encompasses the set of shedder DPI during which contact occurred between shedders and contact birds.

Almost all contacts became infected regardless of shedder DPI, indicating that shedder birds started to become infectious prior to 10 DPI and remained infectious until at least 20 DPI (Fig S3). A much lower proportion of contact birds developed disease symptoms or died when exposed to vaccinated shedders. Results were also suggestive that shedder DPI at exposure may have some influence on disease progression in contact birds: more contact birds developed disease symptoms when exposed to vaccinated shedders from 14 shedder DPI onwards, and more contact birds died after contact with sham-vaccinated shedders at later shedder DPI.

**Fig S3. Impact of shedder vaccination status and days post-infection on contact bird infection, disease symptoms and mortality.** Contacts positive for virus in qPCR from samples taken at 14 DPC were classified as infected. “Diseased” individuals showed visible symptoms (peripheral nerve enlargement and/or tumours) at necropsy, 8 weeks post contact or upon death. “Dead” individuals died due to MD prior to the end of the 8-week experimental period. HVT = contacts exposed to vaccinated shedders; PBS = contacts exposed to sham-vaccinated shedders. The two replicates were pooled for this figure.

**(iii) Determination of appropriate sampling time post-contact for measuring FVL in contact birds.** In the main study, FVL was used as a means to establish successful MDV transmission from shedder to contact birds and to gain insights into the potential mechanisms underlying vaccine effects. To avoid confounding between shedder-contact bird and contact-contact bird transmission, FVL samples of contact birds needed to be taken prior to the onset of contact-contact bird transmission.

Two replicates of experiments carried out with unvaccinated shedders of 4 different ADOL inbred chicken lines (MD-resistant Line 6 and Line 15.N-21, and MD-susceptible Line 7 and Line 15.P-19) revealed that 15I_5_ x 7_1_ F_1_ strain contact bird FVL was low and often below the level of detection at 7 DPC, but significantly higher and above the level of detection at 14 DPC (Fig S4). Hence, for the main experiment FVL in contact birds was measured at 14 DPC, as this provided a reliable indicator of successful transmission between shedder and contact birds, and because 14 DPC lies within the expected period prior to the onset of contact-contact bird transmission and presence of virus in feathers (given an assumed latency period of 7 days^2^).

**Fig S4. Effect of number of days post-contact on contact bird feather viral load.** Histogram of contact bird FVL from qPCR at 7 (red bars) and 14 (blue bars) days post-contact with unvaccinated infectious shedders (2 replicates and all 4 shedder chicken lines combined). A value of -5 indicates negative for MDV, i.e. values were below the level of detection by standard qPCR.
