## Supplementary figures and images for "Pathogen transmission from vaccinated hosts can cause dose-dependent reduction in virulence"

### Fig S1

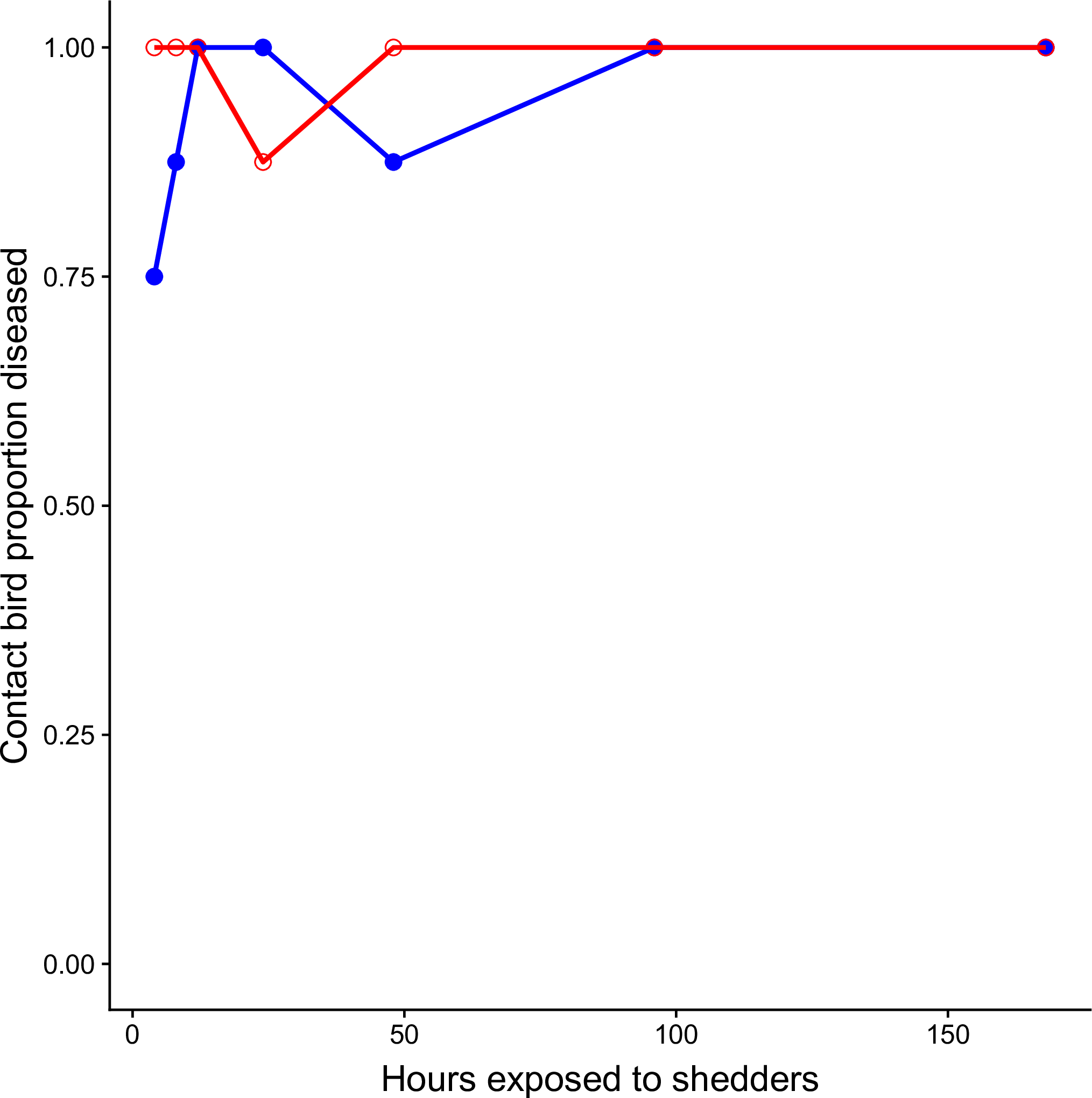

### Fig S2

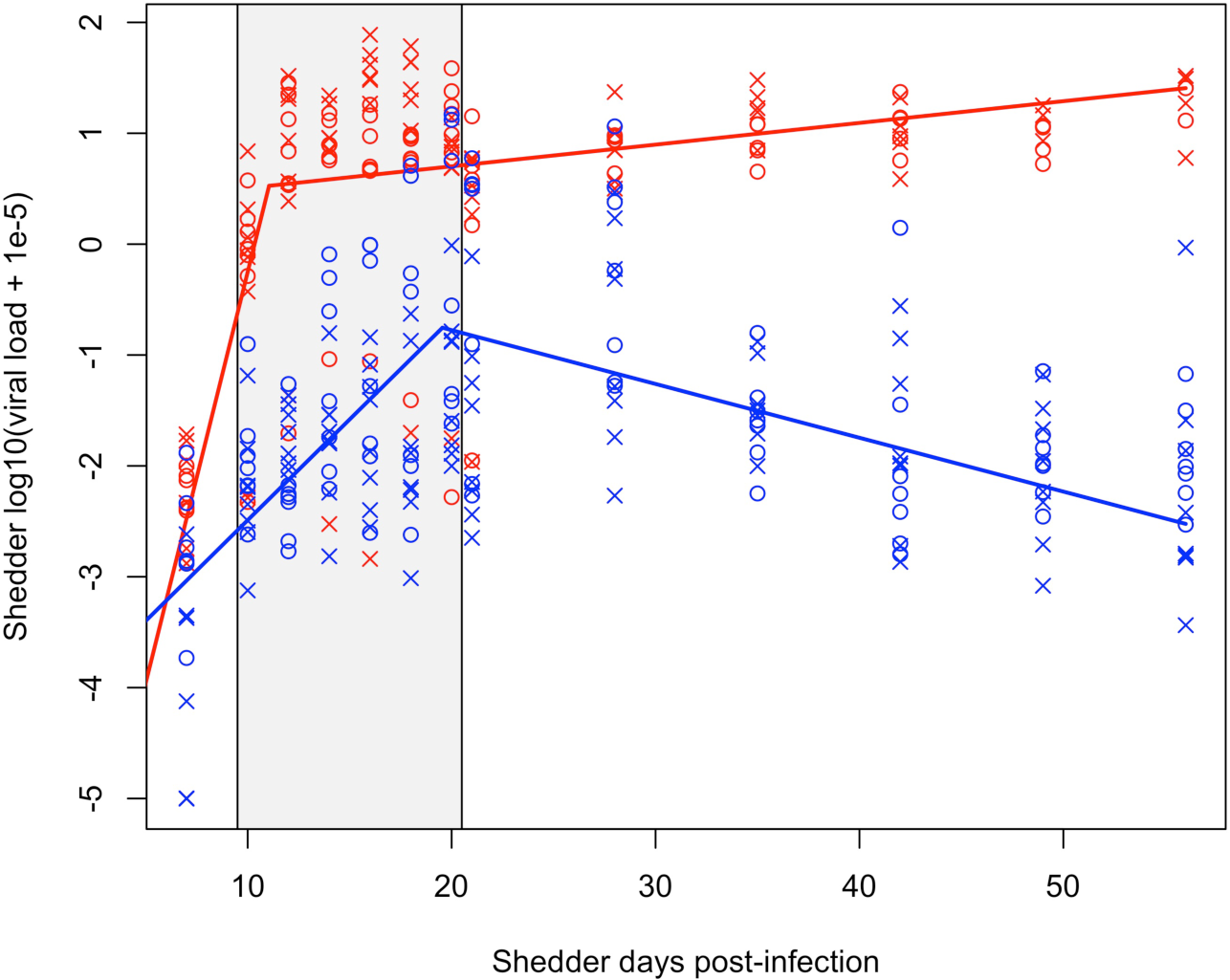

### Fig S3

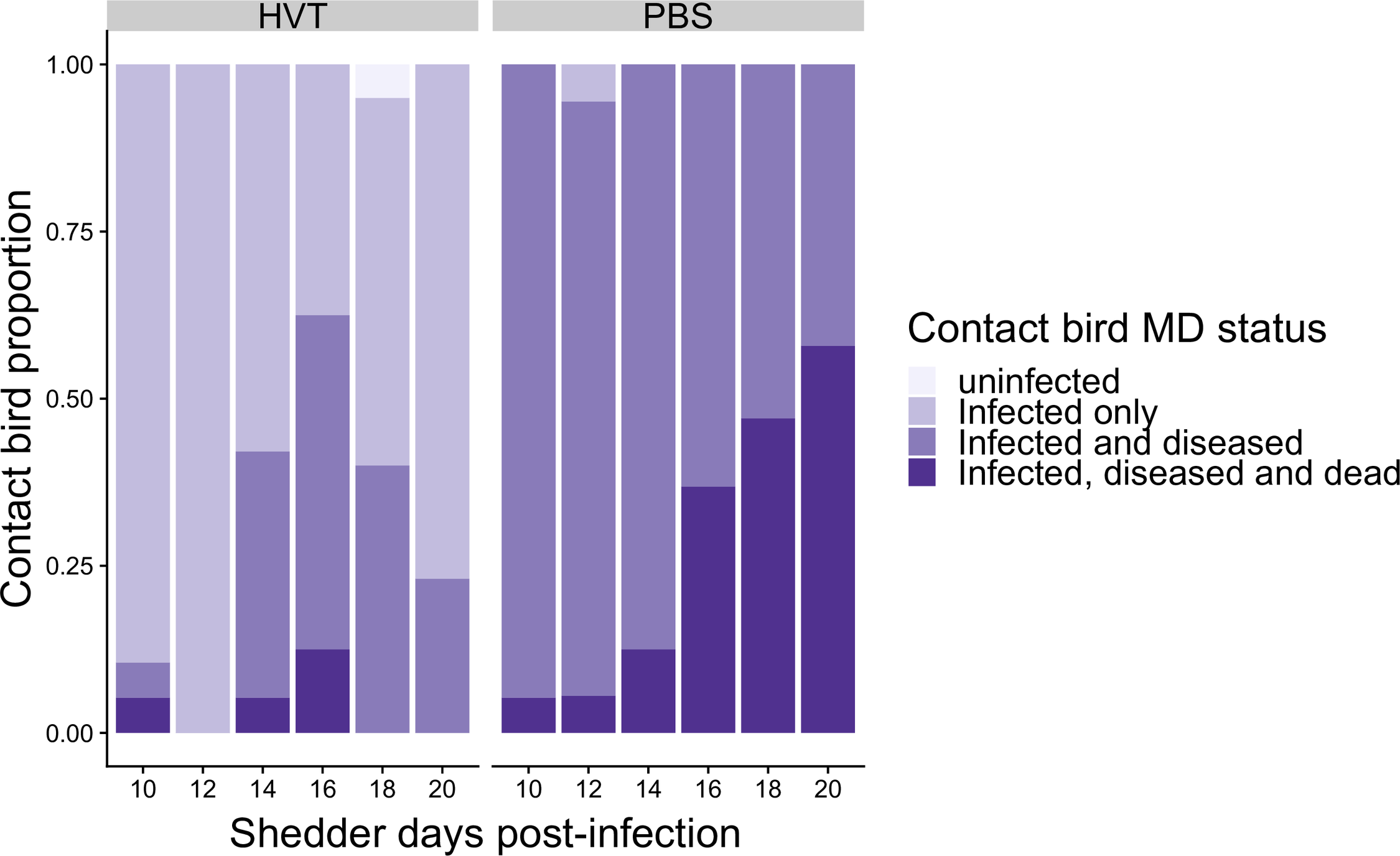

### Fig S4

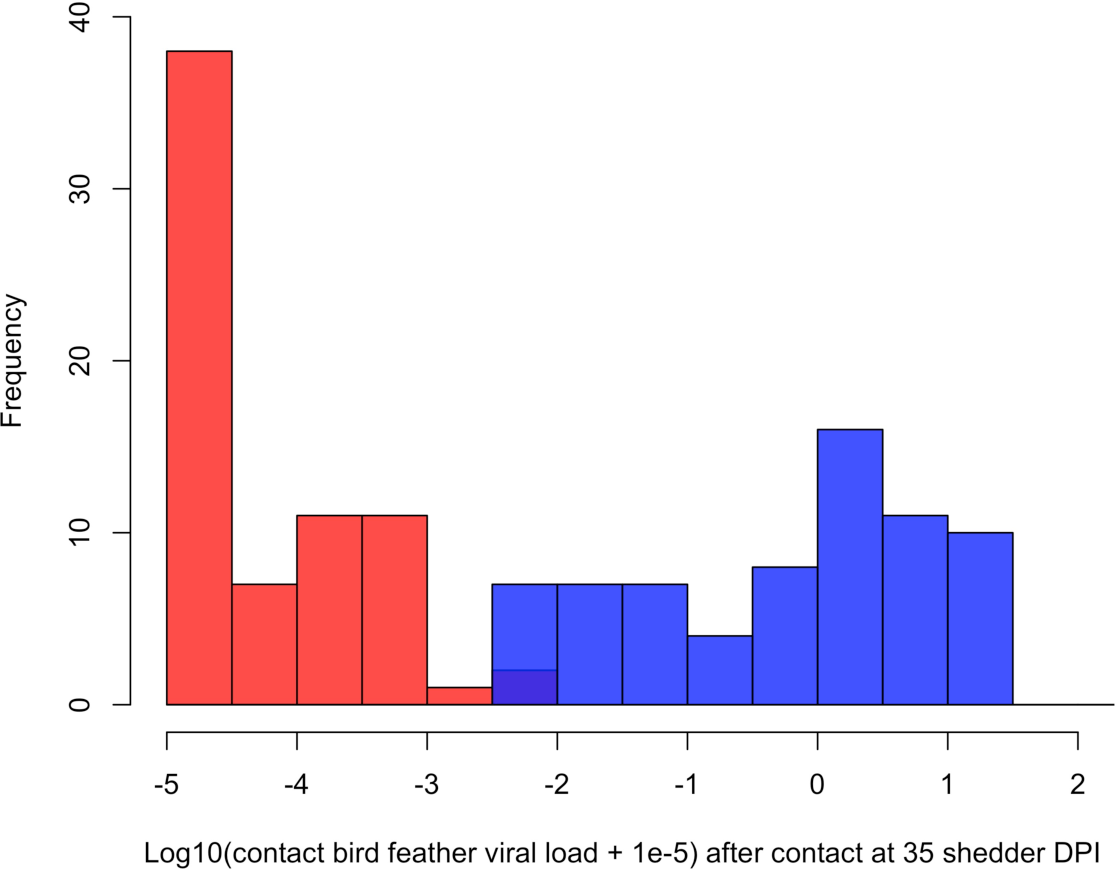
